# Evidence of virulence and antimicrobial resistance in *Streptococcus pneumoniae* serotype 16F lineages

**DOI:** 10.1101/2023.08.25.554804

**Authors:** Jolynne Mokaya, Kate C Mellor, Gemma G R Murray, Akuzike Kalizang’oma, Cebile Lekhuleni, Heather J Zar, Mark P. Nicol, Lesley McGee, Stephen D Bentley, Stephanie W Lo, Felix Dube, The Global Pneumococcal Sequencing Consortium

## Abstract

**Introduction:** Due to the emergence of non-vaccine serotypes in vaccinated populations, *Streptococcus pneumoniae* remains a major global health challenge despite advances in vaccine development. Serotype 16F is among the predominant non-vaccine serotypes identified among vaccinated infants in South Africa (SA).

**Aim:** To characterise lineages and antimicrobial resistance in 16F isolates obtained from South Africa and placed the local findings in a global context.

**Methodology:** We analysed 10923 *S. pneumoniae* carriage isolates obtained from infants recruited as part of a broader SA birth cohort. We inferred serotype, resistance profile for penicillin, chloramphenicol, cotrimoxazole, erythromycin and tetracycline, and Global Pneumococcal Sequence Clusters (GPSCs) from genomic data. To ensure global representation, we also included *S. pneumoniae* carriage and disease isolates from the Global Pneumococcal Sequencing (GPS) project database (n=19,607, collected from 49 countries across five continents, years covered (1995 - 2018), accessed on 17^th^ March 2022).

**Results:** Nine percent (934/10923) of isolates obtained from infants in the Drakenstein community in SA and 2% (419/19607) of genomes in the GPS dataset were serotype 16F. Serotype 16F isolates were from 28 different lineages of *S. pneumoniae,* with GPSC33 and GPSC46 having the highest proportion of serotype 16F isolates at 26% (346/1353) and 53% (716/1353), respectively. Serotype 16F isolates were identified globally, however, most isolates were collected from Africa. GPSC33 was associated with carriage [OR (95% CI) 0.24 (0.09 – 0.66); p=0.003], while GPSC46 was associated with disease [OR (95% CI) 19.9 (2.56 – 906.50); p=0.0004]. 10% (37/346) and 15% (53/346) of isolates within GPSC33 had genes associated with resistance to penicillin and co-trimoxazole, respectively, and 18% (128/716) of isolates within GPSC46 had genes associated with resistance to co-trimoxazole. Resistant isolates formed genetic clusters which may suggest emerging resistant lineages.

**Discussion:** Serotype 16F lineages are common in Southern Africa. Some of these lineages are associated with disease, and resistance to penicillin and cotrimoxazole. We recommend continuous genomic surveillance to determine long term impact of serotype 16F lineages on vaccine efficacy and antimicrobial therapy globally. Investing in vaccine strategies that offer protection over a wide range of serotypes/lineages remains essential.

**DATA SUMMARY:** The sequencing reads for the genomes analysed have been deposited in the European Nucleotide Archive and the accession numbers for each isolate are listed in **Supplementary Table1**. Phylogenetic tree of serotype 16F pneumococcal genomes and associated metadata are available for download and visualisation on the Microreact website: Phylogenies of seotype 16F, GPSC33 and GPSC46 are available on the Microreact <u>serotype-16F</u>, <u>GPSC33</u> and <u>GPSC46</u>, respectively.

**IMPACT STATEMENT:** This study shows that serotype 16F lineages are predominant in Southern Africa and are associated with disease and antimicrobial resistance. Although serotype 16F has been included in the newer formulation of the upcoming vaccine formulations of PCV21 and IVT-25, continuous surveillance to determine long term impact of serotype 16F lineages on vaccines and antimicrobial therapy remains essential.

## INTRODUCTION

Childhood morbidity and mortality caused by *Streptococcus pneumoniae* (*S. pneumoniae*) remains a major global health challenge despite advances in antimicrobial treatment and vaccine development. In 2015, 294,000 HIV-uninfected infants and 23,300 HIV-infected infants were estimated to die from pneumococcal disease globally (1). Pneumococcal conjugate vaccines are the current recommended formulation for children and they target different serotypes that cause most invasive pneumococcal disease (IPD) (2). However, pneumococcal disease persists, in part due to the increase of disease caused by *S. pneumoniae* expressing non-vaccine serotypes, a phenomenon called serotype replacement (3). In serotype replacement, the reduction of vaccine serotypes by PCV or/and antimicrobials creates room for the expansion of non-vaccine serotypes, which can occur between or within lineages (3,4). These non-vaccine serotypes are also sometimes resistant to antimicrobial treatments, leading to an increased risk to public health.

The capsule of *S. pneumoniae* has been classified into 104 serotypes based on the reaction of a set of antisera against the capsular antigen (5). While serotype classification helps to identify virulent strains and subsequently inform vaccine development, it provides little information about the genetic shifts in *S. pneumoniae* strains given that the *cps* locus (which encodes for the capsular polysaccharides) only accounts for 0.2% of the genome (6). Multilocus sequence typing (MLST) was previously the gold standard for characterising bacterial isolates, based on the genetic sequences of seven housekeeping genes, however, recombination in some of these genes and limited resolution inhibits its utility to infer relationships between strains (7,8). Whole genome analysis offers the opportunity to classify strains that share an evolutionary history into lineages allowing for inference of relationships between strains across a species (7). Furthermore, housekeeping genes represent a very small proportion of the genome, and are not necessarily representative of the relationship across the rest of the genome due to recombination within these genes (7). The Global Pneumococcal Sequencing project (GPS), whose aim is to provide an international understanding of *S. pneumoniae* population structure and its impact on vaccine and treatment strategies, has therefore classified *S. pneumoniae* strains into > 900 lineages (also known as Global Pneumococcal Sequencing Clusters [GPSCs]) and each of these lineages can express a single or multiple serotypes (4,7,9).

In South Africa, where PCV13 is part of the expanded programme on immunization (EPI), serotype 16F has been reported to be among the predominant non-vaccine serotypes (NVT) among fully vaccinated infants (10) and has been shown to contribute to invasive disease among children below the age of three years after the introduction of PCV13 (4). Using whole genome sequencing, we characterised the major *S. pneumoniae* lineages expressing serotype 16F in isolates from children in the Western Cape of South Africa and assess the public health relevance of serotype 16F lineages by describing their association with disease and antimicrobial susceptibility.

## METHODS

### Study setting

We analysed 10923 *S. pneumoniae* carriage isolates obtained from 1020 of 1143 infants enrolled from May 2012 – September 2015, to a population-based, longitudinal prospective birth-cohort study [the Drakenstein Child Health Study (DCHS)] in the Western Cape Province in South Africa (11). This study took place at two primary healthcare clinics located two kilometres apart, i.e., TC Newman and Mbekweni. All infants received routine immunisation including PCV13 as part of the national immunisation programme, administered at six weeks, 14 weeks, and nine months of age through a 2+1 vaccine schedule. Nasopharyngeal (NP) swab collection was done every two weeks for the first year of life as well as at six, 12, 18 and 24 months, and whenever infants presented with pneumonia or lower respiratory tract infection (LRTI). Details on sample collection, transportation, culture, and storage have been described previously (10). For global context, we included *S. pneumoniae* isolates of serotype 16F and lineages of interest included in the GPS project database (n=19607) from 49 countries across five continents, years covered (1995 - 2018) (last accessed on 17^th^ March 2022) (12).

### Definition of carriage and disease, and vaccine and non-vaccine serotypes

We define carriage isolates as those collected from healthy individuals and disease isolates those from sputum or other specimen (i.e., blood, joint fluid, aspirates, etc) from individuals with disease (including septicaemia, bacteraemia, pneumonia, cellulitis, meningitis, otitis media, bronchitis, osteomyelitis, septic arthritis, sinusitis, empyema, abscess, surgical site infection, sepsis, encephalitis, lower respiratory tract infection, conjunctivitis, bronchitis, peritonitis, pericarditis). In this manuscript, vaccine serotypes are those included in PCV13, PCV15 and/or PCV20, **Table 1**.

**Table 1:** Serotypes included in Pneumococcal Conjugate Vaccines (PCV)

| PCV Formulation | PCV-7<br>(2000) | PCV-10<br>(2009) | PCV-13<br>(2010) | PCV-15<br>(2019) | PCV-20*<br>(2023) |
| --- | --- | --- | --- | --- | --- |
| **Serotypes included in the vaccine** | 4 | 4 | 4 | 4 | 4 |
|  | 6B | 6B | 6B | 6B | 6B |
|  | 9V | 9V | 9V | 9V | 9V |
|  | 14 | 14 | 14 | 14 | 14 |
|  | 18C | 18C | 18C | 18C | 18C |
|  | 19F | 19F | 19F | 19F | 19F |
|  | 23F | 23F | 23F | 23F | 23F |
|  |  | 1 | 1 | 1 | 1 |
|  |  | 5 | 5 | 5 | 5 |
|  |  | 7F | 7F | 7F | 7F |
|  |  |  | 3 | 3 | 3 |
|  |  |  | 6A | 6A | 6A |
|  |  |  | 19A | 19A | 19A |
|  |  |  |  | 22F | 22F |
|  |  |  |  | 33F | 33F |
|  |  |  |  |  | 8 |
|  |  |  |  |  | 10A |
|  |  |  |  |  | 11A |
|  |  |  |  |  | 12F |
|  |  |  |  |  | 15B |

\*PCV20 (Pevnar20)

### Sequencing and bioinformatics analysis

Single colony picks of presumptive *S. pneumoniae* were inoculated onto Columbia blood agar base with 2% agar, 5% horse blood (BA) plates and incubated at 37^0^C in 5% CO_2_ overnight. DNA was extracted and quantified as described previously (10). DNA was sequenced on an Illumina HiSeq platform at Wellcome Sanger Institute, generating ≥100bp paired-end reads. The reads were assessed for quality, assembled, annotated, and mapped as previously described (13). We inferred serotype using seroBA v1.0.2 and resistance profiles for amoxicillin, cefoxitin, ceftazidime, penicillin, chloramphenicol, clindamycin, cotrimoxazole, doxycycline, erythromycin, levofloxacin, meropenem, rifampicin, tetracycline and vancomycin from the genomic data using the CDC antimicrobial detection tool (14). Multidrug resistance was defined as predicted resistance to ≥03 antibiotic classes. PopPUNK v.1.1.6 was used to assign GPSC to the genomes (15). We mapped reads to GPSC-specific reference [GPSC 46 (accession number ERS628712) and GPSC33 accession number ERS566825)] sequences using BWA v 0.7.17 to create an alignment and then assessed for recombination within each GPSC using Gubbins v2.4.1. Phylogenetic trees for recombination-stripped alignments for each GPSC were generated using RAxML v 8.2.8. To investigate the emergence of serotype 16F lineages, we generated time trees using BactDating with a mixed gamma, relaxed clock model and visualised these trees using FigTree v.1.4.4 (16). Using Gubbins output, we calculated and compared recombination rates of different lineages as follows: sum of the recombination base substitutions across all branches divided by the sum of the point mutations across all branches.

### Descriptive and statistical analysis

We described the proportion of isolates which were serotype 16F using the Drakenstein and GPS datasets. We reported the total number of isolates and serotype 16F isolates within each lineage. To determine the association between serotype 16F lineages with carriage and disease, we only analysed isolates from South Africa to limit heterogeneity - different countries employ varying methods in surveillance, data analysis and data collection approaches. We also only included isolates from children <7 years as pneumococcal disease is most common in this population. For each lineage, we compared the proportions of pneumococcal disease and carriage isolates to those of other lineages, reporting an odds ratio with 95% confidence interval (CI) by Fisher’s exact test. We calculated the frequency of antimicrobial resistance (AMR) for different classes of antimicrobials in each GPSC as follows: (total number of isolates with predicted antimicrobial resistance within a specific lineage/total number of isolates within that lineage) multiplied by 100.

## RESULTS

Characteristics of isolates obtained from infants recruited from the Drakenstein community and isolates obtained from the GPS project are summarised in **Table 2**.

**Table 2:** Characteristics of *Streptococcus pneumoniae* obtained from the Drakenstein community and GPS dataset.

| n (%) |  |  |
| --- | --- | --- |
|  | Drakenstein Community | GPS |
| Total number of isolates | 10923 (100) | 19607 (100) |
| **Clinical manifestation** |  |  |
| Carriage | 10626 (97.3) | 6345 (32.4) |
| Disease<sup>#</sup> | 297 (2.7) | 12435 (63.4) |
| Other<sup>*</sup> | - | 827 (4.2) |

<sup>#</sup> Isolates obtained from sputum collected at the time of lower respiratory tract infection from children participating in the Drakenstein Child Health Study were classified as disease isolates. (-) indicates no data. \* This includes isolates labelled as Dis-Car, UKWN or #N/A in the GPS database.

### Proportion of serotype 16F

A total of 10923 *S. pneumoniae* genomes, comprising 61 serotypes, were isolated from nasopharyngeal swabs taken from the 1020 infants enrolled in the DCHS (**Figure 1**). Serotype 16F was the most common serotype, accounting for 9% (934/10923) of isolates (**Figure 2a**). In the GPS dataset, there were 19607 *S. pneumoniae* genomes, comprising 98 serotypes and serotype 16F had a prevalence of 2% (419/19607) (**Figure 2b**).

**Figure 1:**
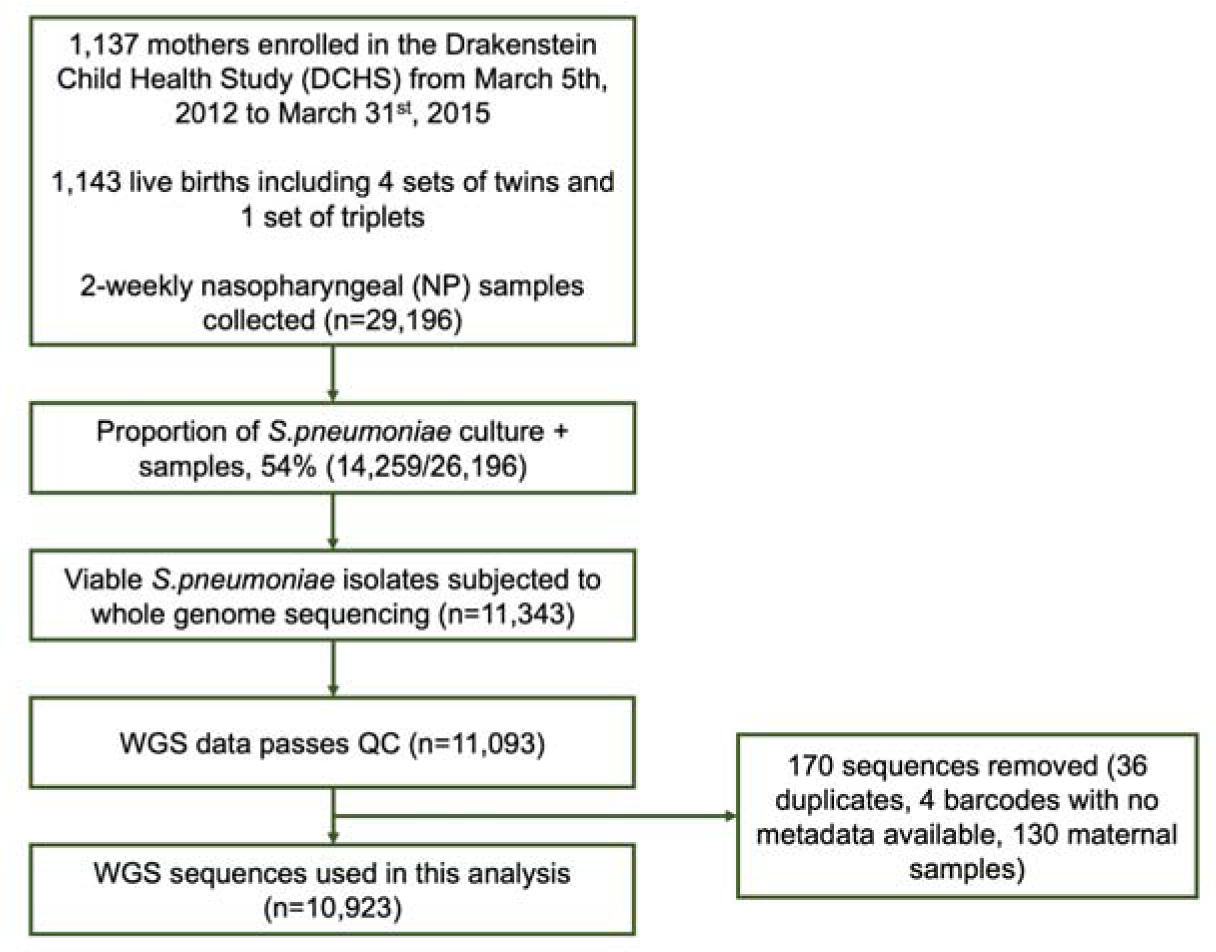
Flow chart of how samples and sequences were included from the Drakenstein community.

**Figure 2:**
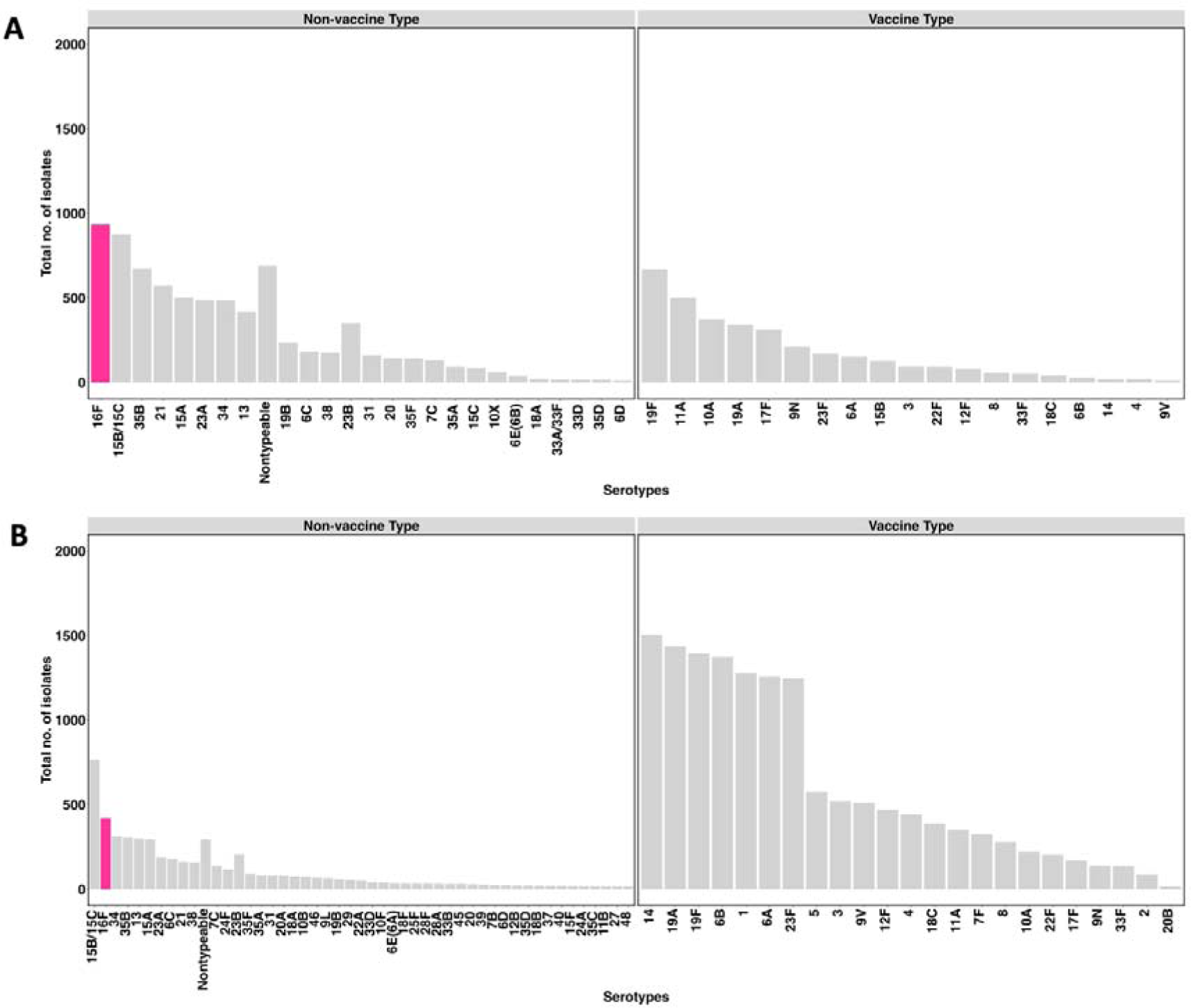
Distribution of *Streptococcus pneumoniae* serotypes within the (A) Drakenstein community and (B) GPS datasets. Bars indicate the number of isolates of each serotype. The pink bar represents serotype 16F and other serotypes shown with grey bars. Serotypes were classified as non-vaccine type (not included in pneumococcal vaccines (PCV)) and vaccine type (included in PCV13, PCV15 and/or PCV20: i.e., 1, 2, 3, 4, 5, 6A/B, 7F, 8, 9N/V, 10A, 11A, 12F, 14, 15B, 17F, 18C, 19A/F, 20B, 22F, 23F, 33F). Serotypes with less than 10 isolates were not shown in these bar plots.

### Global distribution and lineages associated with serotype 16F

We considered the overall geographical distribution of 1353 serotype 16F isolates from the DK and GPS datasets. Serotype 16F was identified in six continents, with the majority of 16F isolates collected from Africa [92% (1248/1353)]. The proportion of serotype 16F isolates from other continents were as follows: 2% (59/1353) were from Asia, 1.8% (25/1353) from North America, 1.4% (19/1353) from South America and 0.15% (2/1353) from Europe, (**Figure 3**). Serotype 16F was present in 28 distinct lineages, with GPSC33 and GPSC46 having the highest proportion of 16F strains, at 26% (346/1353) and 53% (716/1353) respectively (**Figure 3**). Geographical structure was observed within the African continent with GPSC103 and GPSC274 more commonly detected in West Africa, GPSC46, GPSC47, and GPSC207 in Southern Africa, GPSC268 in East Africa, and GPSC33 and GPSC114 on the South and East coast of Africa. In North America, the predominant lineages were GPSC135 and GPSC165. GPSC18, GPSC156 and GPSC104 were predominant in Latin America, Europe, and Asia, respectively.

**Figure 3:**
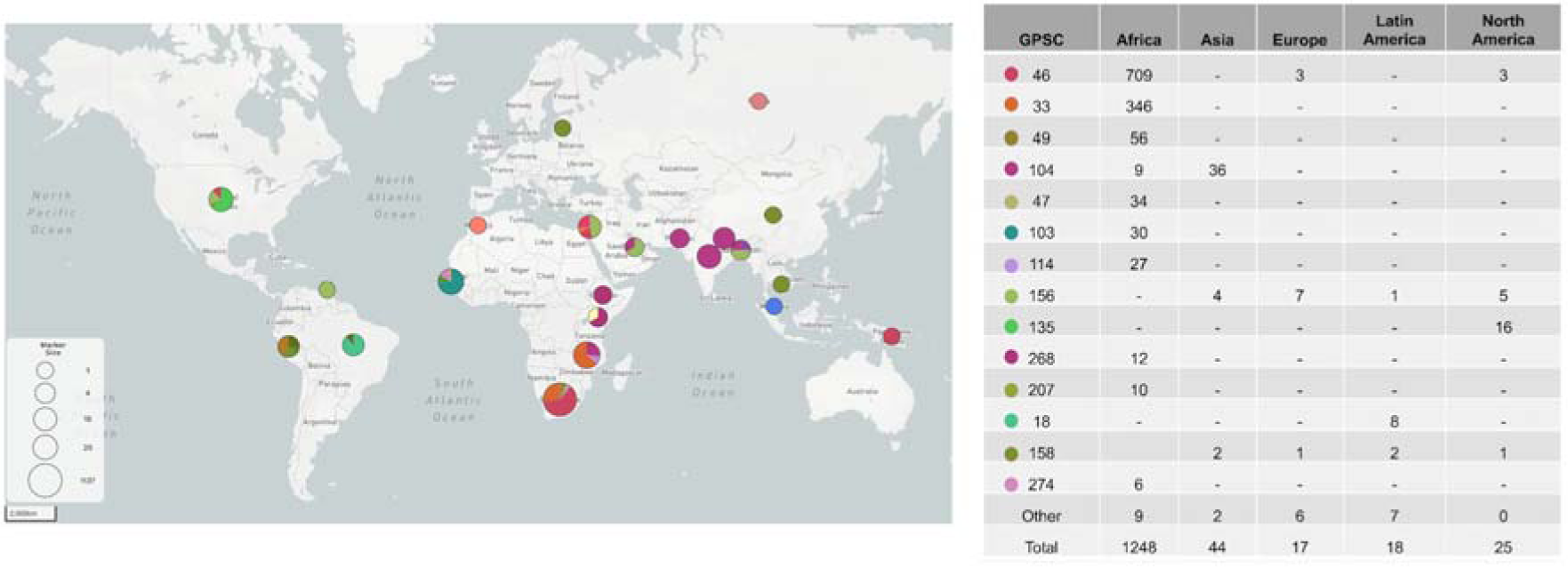
Global distribution of serotype 16F lineages. The pie charts show the proportions of serotype 16F lineages in different countries. GPSC = Global Pneumococcal Sequencing Clusters. Available on the Microreact <u>serotype-16F</u>. Table on the right summarises the number of serotype 16F isolates within each lineage across different continents. GPSCs with less than 5 isolates (GPSC80, 313, 348, 386, 428, 434, 477, 485, 505, 542, 568, 583, 676, 846) are not shown on the table. One pneumococcal isolate from Oceania belonging to GPSC46 was not shown on the table.

### Antimicrobial resistance across serotype 16F lineages

We assessed the presence of AMR genes in serotype 16F lineages (**Figure 4**). 10% (37/346) or 15% (53/346) of isolates within GPSC33 had genes associated with resistance to penicillin or co-trimoxazole, respectively, while 18% (128/716) of isolates within GPSC46 had genes associated with resistance to co-trimoxazole. All isolates (n=30) within the GPSC103 lineage had genes associated with resistance to cotrimoxazole, tetracycline and doxycycline.

**Figure 4:**
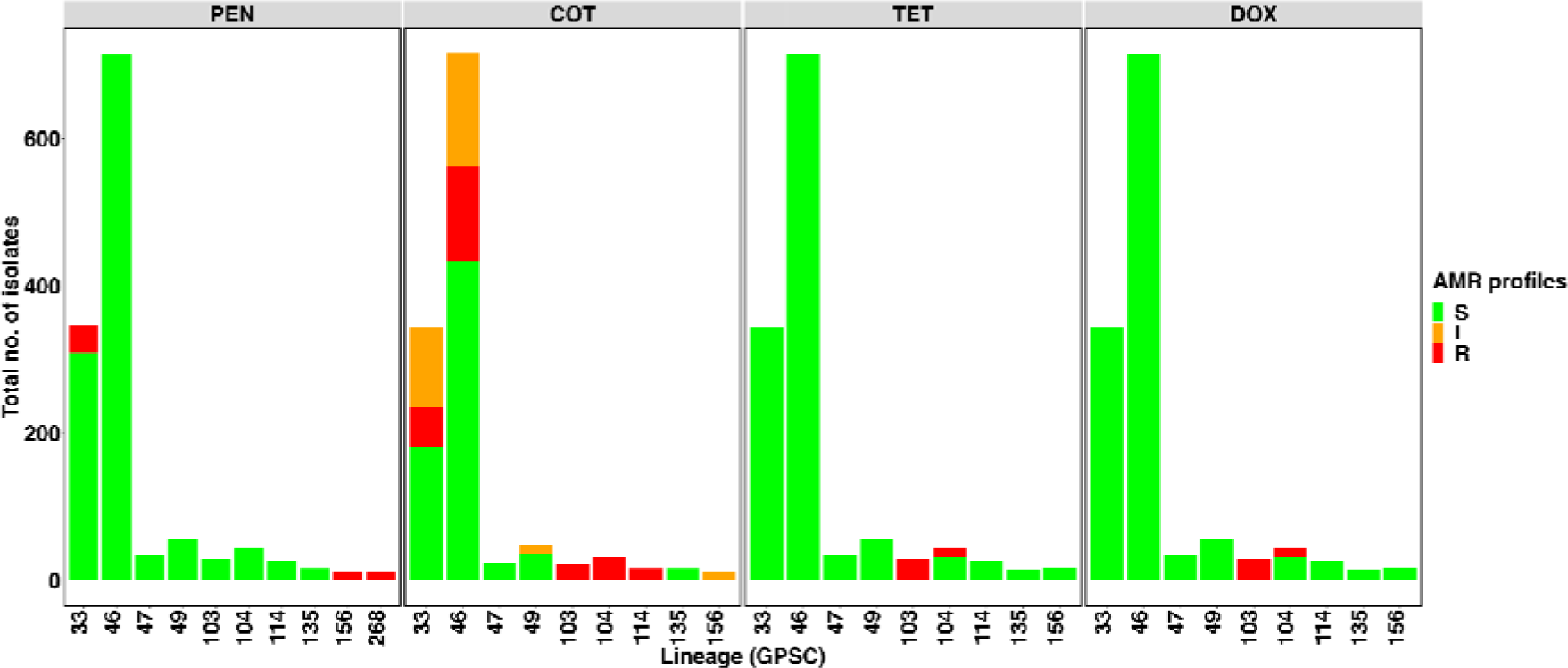
Antimicrobial resistance profiles of serotype 16F lineages. PEN = Penicillin, COT = Cotrimoxazole, TET = Tetracycline and DOX = Doxycycline. S = Susceptible, I = Intermediate resistance and R = Resistant

### Lineage analysis of GPSC33 and GPSC46

We explored the global phylogeny of GPSC33 and GPSC46 given that they are the predominant lineages of serotype 16F (**Figure 5**). GPSC33 lineage consists almost exclusively of (99%, 346/350) of serotype 16F. All isolates within this lineage were from Southern Africa (South Africa: 88% (307/350)] and Malawi: 12% (43/350)]. Similarly serotype 16F predominates (99%, 716/723) in the GPSC46 lineage. Most of isolates within this lineage were from South Africa [98% (711/723)] with few isolates collected from elsewhere [Belarus (n=2), Israel (n=3), Papua New Guinea (n=1), Russia (n=3) and USA (n=3)]. In GPSC33 there were two distinct clusters of isolates with penicillin-resistance determinants and one cluster of isolates cotrimoxazole-resistance determinants; and in GPSC46, there was one cluster of isolates with cotrimoxazole-resistance determinants (**Figure 4**).

**Figure 5:**
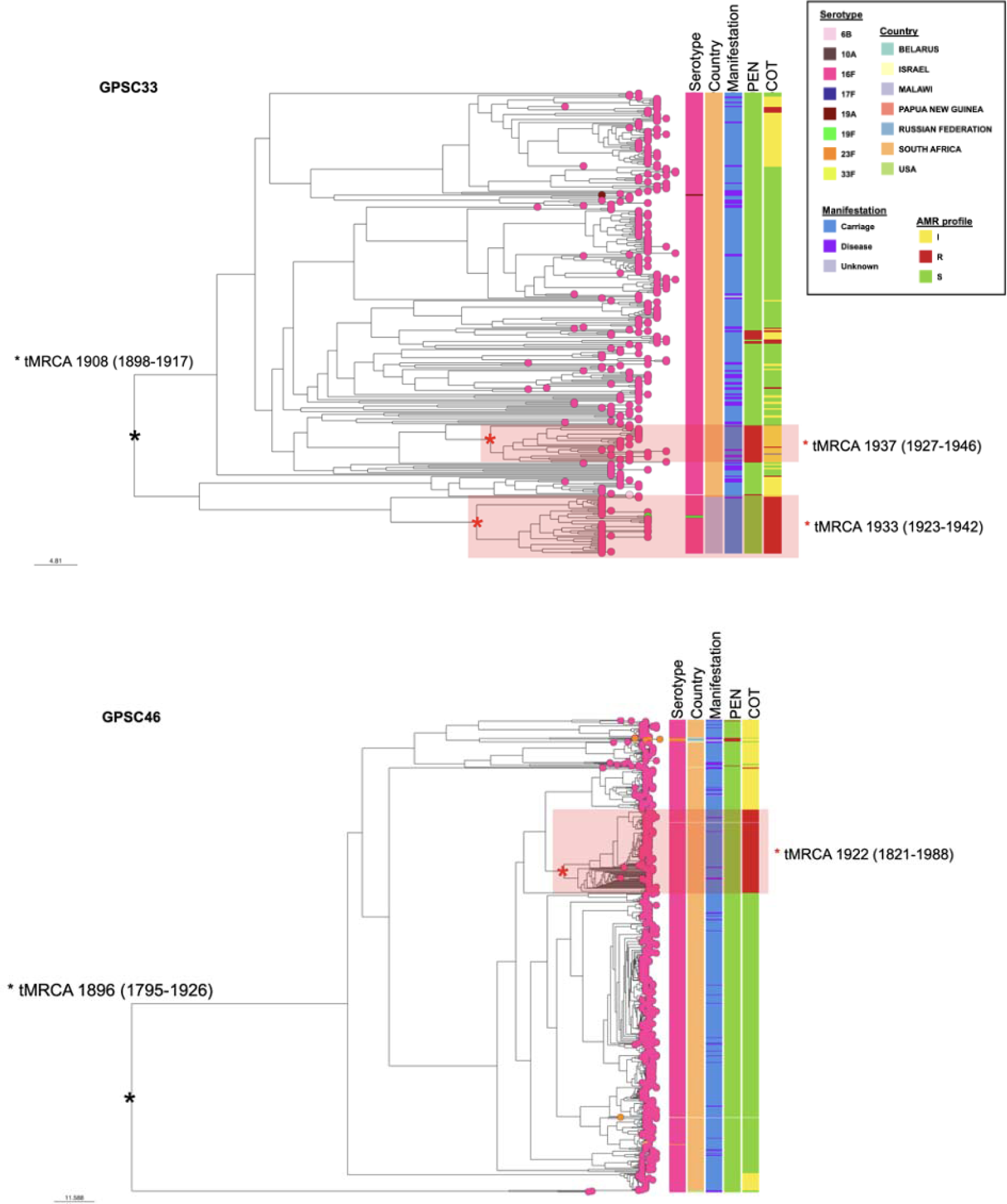
Global phylogeny of GPSC33 and GPSC46. Serotype within each lineage shown on the 1st column on the right of the tree. Countries where the isolates within each lineage were obtained are shown on the 2nd column on the right of the tree. Clinical manifestation of individuals from isolates were obtained from shown on the 3rd column. AMR profile of PEN (penicillin) and COT (cotrimoxazole) shown on the 4th and 5th column on the right of the tree, respectively. I: Intermediate resistance. R: Isolates with resistance. S: Isolates that are susceptible to antimicrobials. Link to access the GPSC33 phylogenetic tree is: <u>GPSC33</u>. Link to access the GPSC46 phylogenetic tree is: <u>GPSC46</u>

Compared to GPSC46, the tMRCA (time to the most recent common ancestor) for GPSC33 is recent; serotype 16F isolates within GPSC33 likely originated in 1908 (95% HPD 1898 - 1917), compared with 1896 (95% HPD 1795 - 1926) for GPSC46. For isolates with resistance formed clusters which suggests emerging resistant lineages, the estimated tMRCA for the cluster with penicillin resistance in GPSC33 is 1937 (95% HPD 1927 - 1946) and 1922 (95% HPD 1821 - 1988) for the cluster with cotrimoxazole resistance in GPSC46. The emergence of AMR clusters might have been driven by treatment exposure given that tMRCA of lineages is between 1920s - 1930s, an estimated time-period when both penicillin and cotrimoxazole were discovered (17,18). We compared the genetic variation through recombination (a process in which exogenous DNA is acquired and incorporated into the genome (19)) of GPSC33 and GPSC46. GPSC33 had a higher recombination ratio (i.e., ratio of the number of recombination events to point mutations on a branch) at 8.2 compared to 4.9 for GPSC46, which could be the reason why there are more clusters with AMR in this lineage.

We compared the association of serotype 16F isolates to other isolates within GPSC33 and GPSC46 to carriage and disease. For this analysis, we only included isolates from South Africa to limit heterogeneity. Compared to other serotypes within GPSC33 and GPSC46 lineages, serotype 16F isolates were associated with carriage in GPSC33 [OR (95% CI) 0.24 (0.09 - 0.66); p=0.003], and with disease in GPSC46 [OR (95% CI) 19.9 (2.56 - 906.50); p=0.0004] (**Table 3**).

**Table 3:**
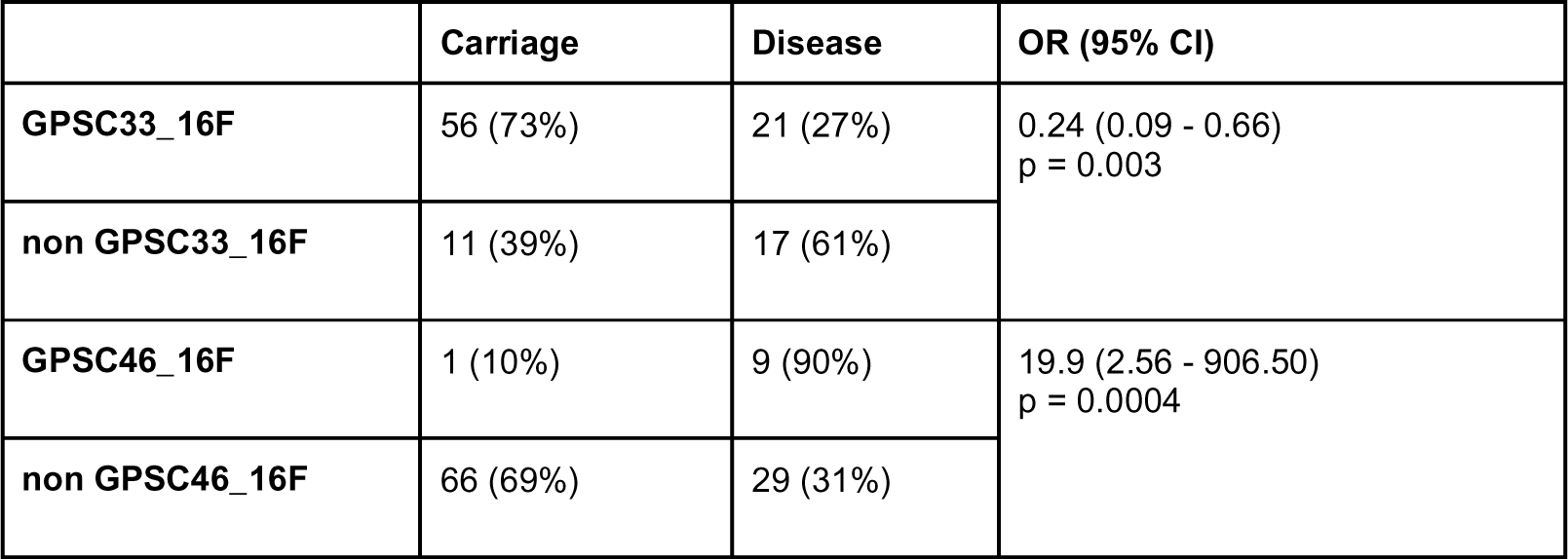
Comparing carriage and disease between GPSC33 and GPSC46.

## DISCUSSION

Serotype replacement from non-vaccine serotypes continues to threaten the effectiveness of current pneumococcal vaccine interventions (11–15). However, pneumococcal surveillance, including use of genomics, has been able to inform vaccine strategies(16). We show that serotype 16F remains a dominant non-vaccine type in both carriage and invasive disease in South Africa. We further contexualised our findings in a global context using the GPS Project Database (7), and we highlight that serotype 16F is prevalent in Africa. Serotype 16F lineages, that include GPSC33 and GPSC46, have been highly successful in the region and are associated with antimicrobial resistance and pneumococcal disease. Together these data highlight that non-vaccine serotype 16F has become increasingly important in South Africa, therefore, further surveillance is warranted to inform vaccine policy and potentially expand vaccine valency.

We show that serotype 16F was the dominant serotype in our longitudinal carriage cohort, similar to other pneumococcal carriage surveillance findings from South Africa (17). Previous work in the region described significant decreases in PCV7 serotypes (19F, 6B, 23F and 14) and an increase in non-vaccine serotypes (16F, 34, 35B and 11A) among children less than two □years of age colonized with the pneumococcus (17). Other regions including West Africa (18) and Middle East (19) have described the predominance of serotype 16F in carriage and disease among children and adults following pneumococcal vaccine rollout. Here we show that lineage GPSC33 was the dominant lineage in our cohort, and the success of the lineage has been described as mostly prevalent on the African continent (18). Longitudinal colonisation studies have shown a typically high number of single nucleotide polymorphisms among 16F isolates of the same GPSCs carried over multiple sampling visits, suggesting divergent 16F strains may emerge over the course of carriage due to homologous recombination (20). Further work is required to understand the genotypic and phenotypic traits of successful lineages such as GPSC33.

Antimicrobial selection and horizontal gene transfer could potentially facilitate the expansion of resistant serotype 16F lineages in South Africa. We identified cotrimoxazole-resistance sub-lineages in both GPSC33 and GPSC46. Although cotrimoxazole is not directly used to treat diseases caused by *S. pneumoniae*, it is widely used as a prophylaxis for all infants who are HIV-exposed but uninfected (21), and adults living with HIV. Antimicrobial pressure due to cotrimoxazole use has therefore been suggested to contribute to the resistance patterns observed in the region, therefore, further use of the antimicrobial may facilitate the expansion of cotrimoxazole resistant GPSC33 and GPSC46 sub-lineages.

Although we established that GPSC33 and GPSC46 lineages were generally susceptible to first-line antimicrobial used to treat pneumonia, such as penicillin, we identified penicillin-resistant GPSC33 sub-lineages, highlighting the potential risk of this lineage expanding and limiting antimicrobial treatment options in South Africa. Penicillin resistant serotype 16F have been described in countries such as Japan following post PCV7 rollout (22). Previous work on horizontal gene transfer has shown that serotypes that frequently colonise the human nasopharynx, including serotype 16F, have been shown to acquire penicillin-binding protein gene fragments from *Streptococcus mitis (23)*. The authors highlighted the presence of mosaic *pbp*2x among serotype 16F GPSC33 associated with reduced (23) susceptibility, and modifications at the *pbp*2x gene are known to confer reduced susceptibility to a range of beta-lactam antimicrobial (24). Due to the high rates of recombination seen among 16F isolates in longitudinal carriage and the acquisition of mosaic pbp fragments from other species (23), we hypothesise that serotype 16F may develop higher levels of beta-lactam resistance with subsequent recombination in other important pbp genes (pbp2b and pbp1a).

Serotype 16F lineages are an important cause of invasive disease in the post-PCV13 era, particularly in countries such as South Africa. Our results show that not only is serotype 16F the predominant non-vaccine serotype carried among vaccinated infants, but serotype 16F lineages that include GPSC33 and GPSC46 cause invasive disease. In South Africa, serotype 16F has been associated with the second highest case fatality ratio after serotype 6A (25). Serotype 16F is an important cause of invasive disease in countries such as Ethiopia (26), Denmark (27), and the serotype has increasingly become important among cases of meningitis through serotype replacement in Israel (19). Previous meta-analyses have shown increased case-fatality rates associated with serotype 16F, along with serotypes 3, 6B, 9N, 11A, 19F, and 19A (28). Furthermore, up to seven studies found an increase for serious clinical outcomes attributed to serotype 16F that is not included in the PCV13 formulation (28). It remains unclear why serotype 16F is associated with mortality, but infection among vulnerable populations may be of concern. There is a predominance of non-vaccine serotypes of the pneumococcus in carriage among the HIV infected children in Ghana, particularly serotype 16F (29). Further genomic surveillance will be important to track serotypes and lineages that are associated with carriage, disease, and elevated risk of serious outcomes for vaccine policy making and potentially expanding vaccine valency to include 16F.

Serotype 16F is not included in the current PCV formulations (PCV10/13/15/20) approved for use in children but is included in upcoming formulations of PCV21 and IVT-25. Therefore, continuous surveillance to determine long term impact of serotype 16F lineages on vaccines and antimicrobial therapy remains essential.

## AUTHOR CONTRIBUTIONS

Conceived the study: JM, SWL, FD. Assimilated data: JM, SWL, FD. Analysed the data: JM. Wrote the manuscript: JM. Revised the manuscript: JM, AK, MPN, SWL, FD. All authors have read and approved the manuscript.

